# Loss of motor cortical inputs to the red nucleus after central nervous system disorders in non-human primates

**DOI:** 10.1101/2022.10.04.510771

**Authors:** Simon Borgognon, Eric M. Rouiller

**Affiliations:** Center for the Neural Basis of Cognition, Department of Bioengineering, University of Pittsburgh (PA), USA; Department of Neurosciences and Movement Sciences, Section of Medicine, Faculty of Sciences and Medicine, University of Fribourg, Fribourg, Switzerland; Center for Neuroprosthetics and Brain Mind Institute, École Polytechnique Fédérale de Lausanne (EPFL), Lausanne, Switzerland

## Abstract

The premotor (PM) and primary motor (M1) cortical areas broadcast voluntary motor commands through multiple neuronal pathways, including the corticorubral projection that reaches the red nucleus (RN). However, the respective contribution of M1 and PM to corticorubral projections as well as its plasticity following motor disorders or injuries are not known in non-human primates. Here, we quantified the density and topography of axonal endings of the corticorubral pathway in RN in intact monkeys, as well as in monkeys subjected to either cervical spinal cord injury (SCI), Parkinson’s disease (PD)-like symptoms or primary motor cortex injury (MCI). Twenty adult macaque monkeys were injected with the biotinylated dextran amine (BDA) anterograde tracer either in PM or in M1. We developed a semi-automated algorithm to reliably detect and count axonal boutons within the magnocellular (mRN) and parvocellular (pRN) subdivisions of RN. In intact monkeys, PM and M1 preferentially target the medial part of the ipsilateral pRN, reflecting its somatotopic organization. PM’s projection to the ipsilateral pRN is denser than M1’s, matching previous observations for the corticotectal, corticoreticular, and corticosubthalamic projections (Fregosi et al., 2018, 2019; Borgognon et al., 2020). In all three types of motor disorders, there was a uniform and strong decrease (near loss) of the corticorubral projections from PM and M1. The RN may contribute to functional recovery after SCI, PD and MCI, by reducing direct cortical influence. This reduction possibly privileges direct access to the final output motor system, via emphasis on the direct corticospinal projection.

## Introduction

The multiple motor cortical areas in primates (primary, premotor, supplementary and cingulate motor cortices) broadcast commands through parallel neuronal pathways to generate and coordinate voluntary movements ^1–3^. These routes reach multiple regions of the central nervous system (CNS) including subcortical relays, the brainstem, and the spinal cord. These projections are specifically organized depending on their origin and their targets ^4^. For instance, in intact adult macaques, the premotor cortex (PM) sends denser projections to the subthalamic nucleus ^5^, the superior colliculus ^6^ and the reticular formation ^7^ than the primary motor cortex (M1). After motor disorders, these projections undergo unique regional plasticity that depends on the lesion type, the severity of the symptoms, and the application of a tentative treatment. First, the density of M1-corticobulbar projection increases after spinal cord injury, regardless of the presence of anti-Nogo-A antibody treatment, whereas it reduces in the case of Parkinson’s disease (PD)-like symptoms treated with autologous cellular therapy ^8^. The corticobulbar projection arising from PM reduces after either M1 injury or PD-like symptoms treated with cellular therapy ^8^. Second, following primary motor cortex injury (MCI), the corticotectal projection density from PM to the superior colliculus reduces when treated with anti-Nogo-A antibody, but does not change in absence of treatment ^9^. In case of PD-like symptoms treated with autologous cellular therapy, there is no consistent impact on the corticotectal projections arising from PM or M1 ^9^. Third, the projections from M1 or PM to the subthalamic nucleus re-organize in a severity-of-lesion manner in PD monkeys treated with autologous cellular therapy: reduction of density after low symptoms and increase of density after severe symptoms ^5^. Region-specific plasticity represents therefore a key neural mechanism that needs to be further investigated. Indeed, new findings can illuminate fundamental principles and may lead to the development of new targeted therapeutic avenues.

Among the subcortical structures receiving direct motor cortical inputs, the red nucleus (RN) constitutes a major player in movement and programming function through the rubrospinal and rubro-olivo-cerebellar systems (e.g. for review: ^10^). Located in the ventral midbrain, RN is subdivided between a magnocellular part (mRN) and a parvocellular part (pRN), characterized by larger sparse neurons and medium-sized neurons, respectively. In primates, evolutionary constraints lead to a complete anatomical segregation and functional specialization of RN ^11–13^. Indeed, the rubrospinal tract originates from mRN and the rubro-olivo-cerebellar tract from pRN. As far as the motor cortical projection is concerned, pRN receives denser projections than mRN ^14^. However, the respective contribution of M1 and PM to the corticorubral projections is not known. Furthermore, the axonal endings topography of these corticorubral projections has not been quantified. Finally, although RN plays a role after PD ^15–17^ and after pyramidal lesion ^18,19^, how the corticorubral projections re-organize following motor disorders or injuries has not been established in primates.

What is the topography and density of the corticorubral projections arising from M1 and PM to the RN? What is the impact of a CNS disorder affecting the motor system on these projections? To answer these questions, we injected 20 adult macaques (*Macaca fascicularis*) with the biotinylated dextran amine (BDA) anterograde tracer either in PM or in M1. To quantify the axonal *terminal boutons* and *en passant boutons* within the RN, we developed an algorithm that detects BDA-stained axonal boutons in pRN and mRN, separately. We first compared the axonal boutons density in the bilateral pRN or mRN arising from PM or M1 in healthy monkeys (n=6). In the same monkeys, we quantified the topography of axonal boutons along the dorso-ventral and medio-lateral axes. We next compared the axonal boutons density differences between the healthy group and (1) four M1 or PM injected monkeys after inducing PD-like symptoms; (2) six M1 injected macaques after inducing cervical SCI and; (3) four PM injected macaques subjected to a lesion in the hand area of M1 (primary motor cortex injury, MCI).

## Results

To quantify the corticorubral projections, we unilaterally injected 6 healthy adult macaques with the anterograde tracer BDA in M1 or in PM (**Figure 1a**; see each individual BDA uptake reconstruction in **Supplementary Figure 1**). The identities of the monkeys that received BDA injections in PM were CH, R12 and R13. BDA injections were located in both the ventral (PMv) and dorsal (PMd) aspect of PM in CH and R13, whereas BDA injections covered only PMd in R12. The monkeys injected in M1 were M310, Z182 and M9380. These injections spanned the arm and hand regions of M1 in M310 and Z182. All these 6 monkeys were involved in previously published reports studying the corticosubthalamic ^5^, corticotectal ^6^ and corticoreticular ^7^ projections. In each monkey, we identified the pRN and the mRN based on a series of Nissl-stained histological sections (**Figure 1b**, left panel) and measured the total rostro-caudal extent of each sub-structure. The total rosto-caudal lengths of the RN were 3.75mm (CH), 4.5mm (R12), 4.25mm (R13), 4.0mm (M310), 4.4mm (Z182) and 4.4mm (M9380) (**Figure 1b**, right panel). As expected, the pRN was located more rostrally than the mRN and covered about two third of the total RN. The mRN occupied the caudal portion of the RN and represented about one third of the total RN (**Figure 1b**, right panel, **Supplementary Figure 2**). An adjacent series of BDA-stained histological sections that spanned the RN allowed us to visualize the axonal boutons emitted by the coricorubral axons (**Figure 1c**, **Supplementary Figure 2**). We then co-registered BDA-stained sections with their corresponding Nissl-stained sections. On the co-registered images, we manually defined the pRN and mRN contours and extracted the RN (**Supplementary Figure 3**). We developed an algorithm that detected the intensity, the size and the circularity of conglomerated pixels to quantify the axonal *terminal boutons* and *en passant boutons* (see Methods, **Figure 1c, Supplementary Figure 3**). The axonal boutons sizes were comparable to the ones found in either the reticular formation, the subthalamic nucleus,the superior colliculus, or the vast majority of boutons in the thalamus, but were much smaller than the few giant endings terminating in the thalamus ^20^.Finally, we computed the density of axonal boutons in 2,500μm^2^ sectors to generate a density map for each section (**Figure 1d, step 1**). For each histological section, we divided pRN and mRN into four quadrants: dorso-lateral, dorso-medial, ventral-lateral and ventro-medial (**Figure 1d, step 2**). To compare the density topography across sections, we interpolated (see Methods) the density of each quadrant into equal sizes and plotted the average density across sections (**Figure 1d, step 3**). The density of each section was measured with the area under the curves (AUC) (see Methods, **Figure 1d**).

**Figure 1.**
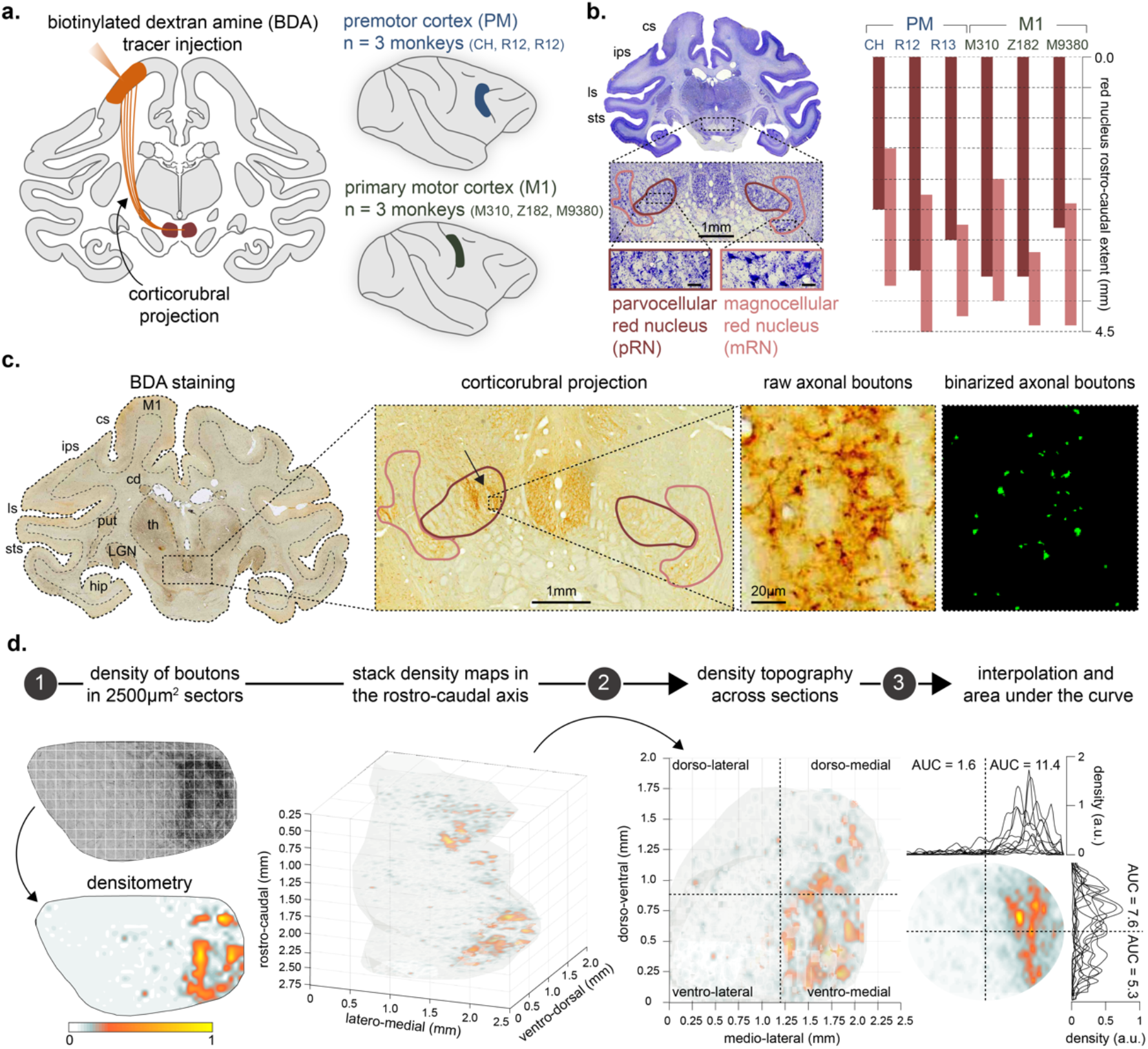
Experimental framework and methodology (illustrated for the 6 intact monkeys). **a.** Left panel: schematic representation of the corticorubral projection in a frontal brain section. We unilaterally injected the anterograde tracer biotinylated dextran amine (BDA) in PM or M1, as schematically represented in the right panel. Three monkeys (CH, R12 and R13) were injected in the premotor cortex (PM) and three other monkeys (M310, Z182 and M9380) in the primary motor cortex (M1). Injection sites reconstructions for each monkey are shown in **Supplementary Figure 1**. **b.** Left panel: representative example of a Nissl-stained histological section showing the segregation between parvocellular red nucleus (pRN, dark red) and magnocellular red nucleus (mRN, light red). Scale bar on the bottom microphotographs = 100μm. Right panel: rostrocaudal extent of the red nucleus for each of the 6 monkeys. More examples of histological sections are shown in **Supplementary Figure 2**. **c.** Semi-automated algorithm to detect axonal boutons (see **Supplementary Figure 3** and Methods). Far left panel: corresponding BDA-stained histological section of **b.** Middle left panel: magnification of the red nucleus showing the corticorubral projection (black arrow). Middle right panel: magnification of axonal boutons located in the ipsilateral pRN. Far right panel: binarized axonal bouton detection based on the intensity, the circularity and the size of conglomerated pixels. **d.** Methodology to compute topography of the corticorubral projections. Step 1: we computed the density of boutons for each histological section. Step 2: we stacked all the sections on the rostro-caudal axis. Step 3: for each histological section, we divided pRN and mRN into four quadrants: dorso-lateral, dorso-medial, ventral-lateral and ventro-medial parts. To compare the density topography across sections, we interpolated (see Methods) the density of each quadrant into equal sizes and plotted the average density across sections. The density of each section was measured with the area under the curves (AUC).

### Intact monkeys

In all intact monkeys (n=6), the ipsilateral pRN represented the preferred corticorubral target regardless the injected motor cortical area (mean across monkeys = 4056 axonal boutons). The second most predominant projection reached the contralateral pRN with an average of 368 axonal boutons across all 6 intact monkeys. The mRN represented only a small proportion of axonal boutons (means were 210 and 76 axonal boutons for the ipsilateral and contralateral mRN, respectively) (**Figure 2a**; p-value < 0.05; Wilcoxon signed rank test). Detailed analyses for each intact monkey separately are shown in **Supplementary Figure 4**. Each monkey showed a comparable distribution of axonal boutons along the rostro-caudal axis (**Supplementary Figure 4a**). We performed the same analysis as shown in **Figure 2a** but for each monkey separately. All intact monkeys showed statistically significant differences between all the paired comparisons possible except for ipsilateral mRN versus contralateral mRN in R12 and R13, and contralateral pRN versus contralateral mRN in M9380 (**Supplementary Figure 4b**, p-values < 0.05, Wilcoxon signed rank test).

**Figure 2.**
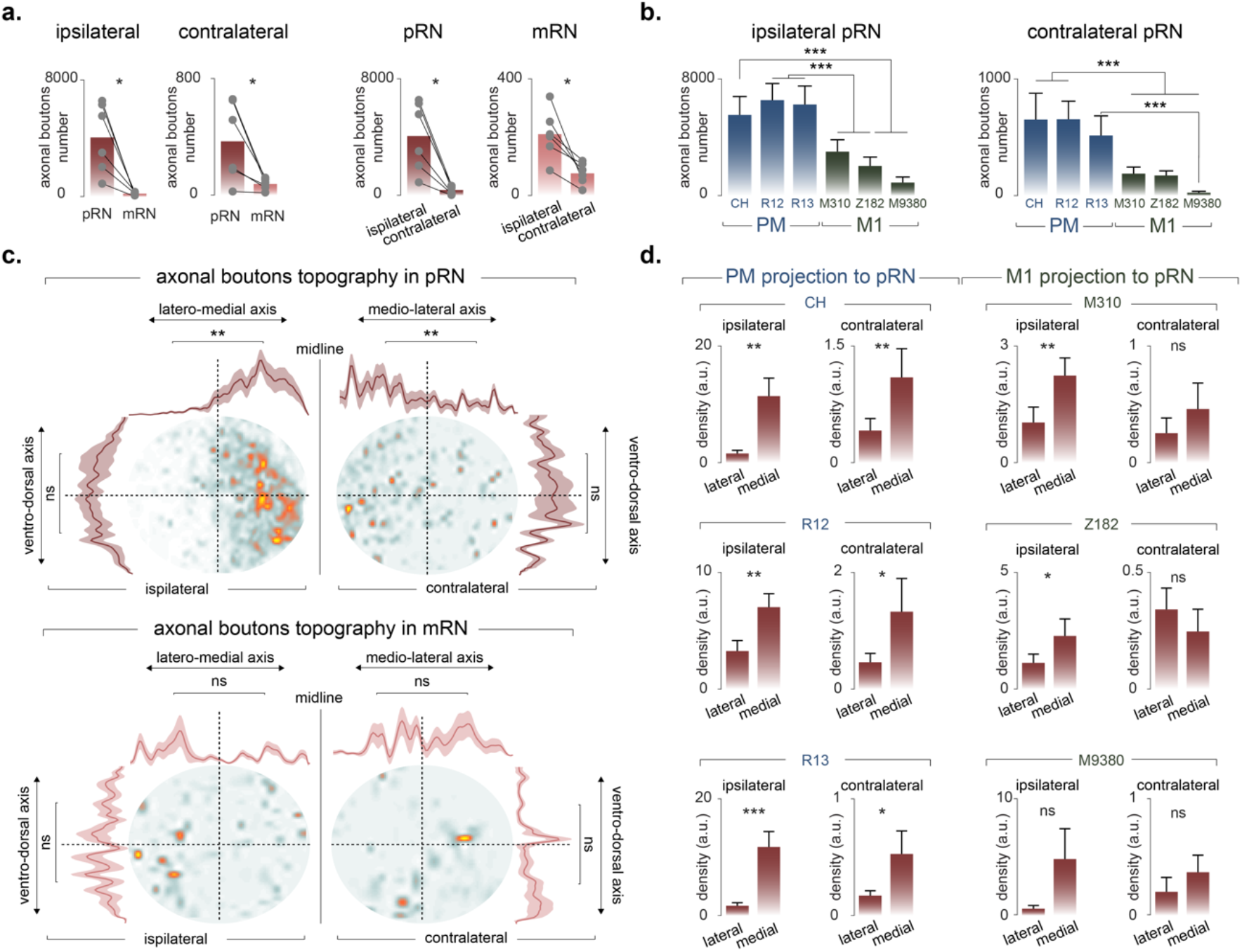
Corticorubral projection density and topography in the 6 intact monkeys. **a.** Bar plots showing the mean axonal boutons number across monkeys (gray dots). Statistically significant differences are indicated with * for p-values ≤ 0.05, Wilcoxon signed rank test **b.** Bar plots showing the means + the standard deviations of axonal boutons number for each individual monkey across bootstrapped histological sections (see Methods). All statistically significances are shown with *** for p-values < 0.00016, residual estimation of the combined distributions and Bonferroni correction for multiple-group comparisons. **c.** Axonal boutons topography in one representative monkey (CH). The curves show the mean ± standard error (dashed areas) across histological sections. The density differences between medial versus lateral and dorsal versus ventral are shown with ** for p-values ≤ 0.01 Wilcoxon signed rank test. “ns” meaning statistically non-significant (p-values > 0.05). **d.** Axonal boutons topography in pRN for each individual monkey. Statistically significant differences are indicated with * for p-values ≤ 0.05, ** for p-values ≤ 0.01, *** for p-values ≤ 0.001, “ns” meaning statistically non-significant (p-values > 0.05) Wilcoxon signed rank test.

Within each injection subgroup (PM or M1), the monkeys yielded comparable numbers of axonal boutons in pRN (statistically nonsignificant; p-values > 0.0083; residual estimation of the combined bootstrapped distributions and Bonferroni correction for multiple-group comparisons, see Methods). However, the number of axonal boutons originating from PM was higher than those from M1 (p-values < 0.0083, residual estimation of the combined distributions and Bonferroni correction for multiple-group comparisons) in all 2 by 2 inter-monkey comparisons, except CH compared to M310 (p-value > 0.0083, residual estimation of the combined bootstrapped distributions and Bonferroni correction for multiple-group comparisons). Likely due to the low numbers of axonal boutons in mRN as compared to pRN, the total numbers of axonal boutons found in mRN showed inconsistency across the two subgroups of intact monkeys (PM versus M1 injections) (**Supplementary Figure 5**).

The projection topography analysis revealed that both PM and M1 preferentially targeted the medial part of the pRN, without a preference along the dorso-ventral axis (**Figure 2c**, top panel). On the other hand, projections to mRN did not show any topographical preference (**Figure 2c**, bottom panel). Quantification of the PM projection topography demonstrated statistically significant increase of density going from lateral to medial in both, the ipsilateral and contralateral pRN (**Figure 2d**, left panel, p-values < 0.05, Wilcoxon signed rank test). As far as the M1 projection topography is concerned, only ipsilateral pRN displayed preferential medial projections (**Figure 2d**, right panel, p-values < 0.05, Wilcoxon signed rank test).

### Monkeys subjected to motor disorders

Given the functional importance of the RN in control of movement, we next aimed at investigating the plasticity of PM and M1 corticorubral projections following a disorder affecting motor functions. We first examined the corticorubral projection in 6 monkeys subjected to SCI, consisting of a cervical hemisection at C7-C8 level, ^21–23^ (**Supplementary Figure 6a**). The BDA injections were performed unilaterally in M1 on the contralesional side. This group was further subdivided into two: a subgroup that received an anti-Nogo-A antibody treatment (n=3, AC, AG and AP) and a control subgroup (n=3, CS, CP and CG) that received a control antibody. All the 6 SCI monkeys exhibited a loss (or dramatic reduction) of M1 originating corticorubral projections (**Figure 3a**; p-values < 0.0071; residual estimation of the combined bootstrapped distributions).

**Figure 3.**
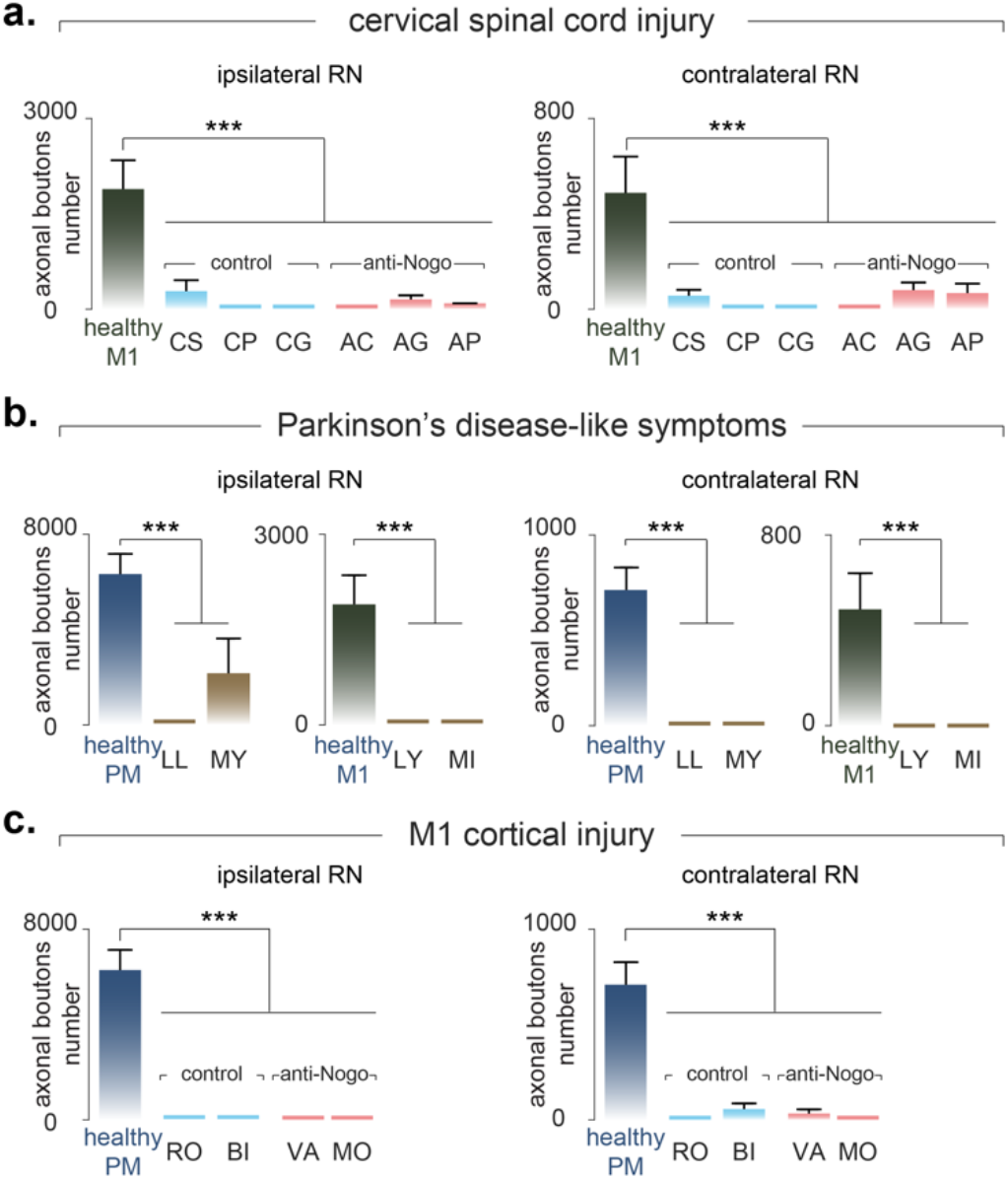
Corticorubral projection plasticity after cervical spinal cord injury, Parkinson’s disease-like symptoms and primary motor cortex (M1) injury. Bar plots showing the means + the standard deviations of axonal boutons numbers across bootstrapped histological sections (see Methods). All statistically significances are shown with *** for p-values < 0.0071 (in **a.**), p-values < 0.016 (in **b.**) and p-values < 0.01 (in **c.**), residual estimation of the combined distributions and Bonferroni correction for multiple-group comparisons. As a result of only few numbers of axonal boutons in the injured monkeys, we pooled the number of axonal boutons of pRN and mRN. The lesion extent as well as the BDA reconstruction uptake are shown in **Supplementary Figure 6. a.** Comparison between intact monkeys injected in M1 (n=3; mean value) and individual monkeys subjected to a unilateral cervical spinal cord injury (n=6). The spinal cord injury group was further subdivided into two groups: a control group (n=3; CS, CP & CG, light blue) that received “sham” treatment and an anti-Nogo-A antibody treated group (n=3; AC, AG & AP; light red) ^21–23^. **b.** Comparison between intact monkeys injected in M1 (n=3; average value) or PM (n=3; average value) and 4 individual monkeys exhibiting Parkinson’s disease-like symptoms (PD) and treated with autologous neural cell ecosystem ANCE ^24,25^. **c.** Comparison between 3 intact monkeys unilaterally injected in PM (average value) and four individual monkeys after a unilateral M1 cortical injury induced with acid ibotenic. The BDA injection was performed in PM homolateral to the injured M1. The M1 cortical injury group was further subdivided into two subgroups: an untreated control subgroup (n=2; RO & BI; light blue) and an anti-Nogo-A antibody treated subgroup (n=2; VA & MO; light red) ^26,27^.

Secondly, we sought to examine the plasticity of the corticorubral projections arising from PM or M1 in a group of monkeys (n=4) experiencing Parkinson’s disease (PD)-like symptoms ^24,25^. Monkeys were administered the neurotoxin 1-methyl-4-phenyl1,2,3,6-tetrahydropyridine (MPTP) to produce symptoms. After reaching a stable plateau of spontaneous functional recovery, all four monkeys received a cellular therapy treatment based on the autologous neural cell ecosystem (ANCE) approach ^24,25^. At the end of the behavioral protocol, two monkeys (LL and MY) were injected with BDA unilaterally in PM (PMd and PMv). The other two monkeys (LY and MI) were injected unilaterally in M1 (hand and arm regions) (**Supplementary Figure 6b**). Similar to the SCI monkeys above, both PD subgroups showed a massive reduction of corticorubral projections originating from PM and M1 (**Figure 3b**; p-values < 0.025; residual estimation of the combined boostrapped distributions).

Finally, we investigated the re-organization of the corticorubral projections originating from PM in M1 cortically injured monkeys (MCI; n=4), all were subjected to a unilateral and permanent lesion of the arm and hand regions of M1 obtained with acid ibotenic injections (**Supplementary Figure 6c**). The four MCI monkeys were subdivided into two subgroups: two monkeys (VA and MO) received anti-Nogo-A antibody treatment and two monkeys (RO and BI) were untreated ^26,27^. All four MCI monkeys were injected with BDA unilaterally in PM, homolateral to the M1 lesion. Surprisingly, the four monkeys also expressed an extensive reduction of corticorubral projections originating from PM (**Figure 3c**; p-values < 0.0125; residual estimation of the combined boostrapped distributions). To sum up, the corticorubral projections arising from PM or M1 were massively reduced regardless of the nature of the motor disorder tested or the respective treatment.

## Discussion

The topography of the corticorubral projections originating from M1 and PM in non-human primates have been previously described in numerous studies ^14,28–32^. Our results are largely consistent with these reports. The corticorubral projections arising from PM and M1 preferentially target the ipsilateral pRN followed by the contralateral pRN, the ipsilateral mRN and finally the contralateral mRN. The originality of the present study consists in the three following points: (i) we reliably quantified the axonal boutons and established a more extended projection topography; (ii) we provided new evidence in intact monkeys that the projection from PM to pRN is denser than that from M1 to pRN (summary in **Figure 4a**); (iii) the corticorubral projections arising from PM and M1 were drastically reduced after motor disorders like SCI, PD and MCI (summary in **Figure 4b**).

**Figure 4.**
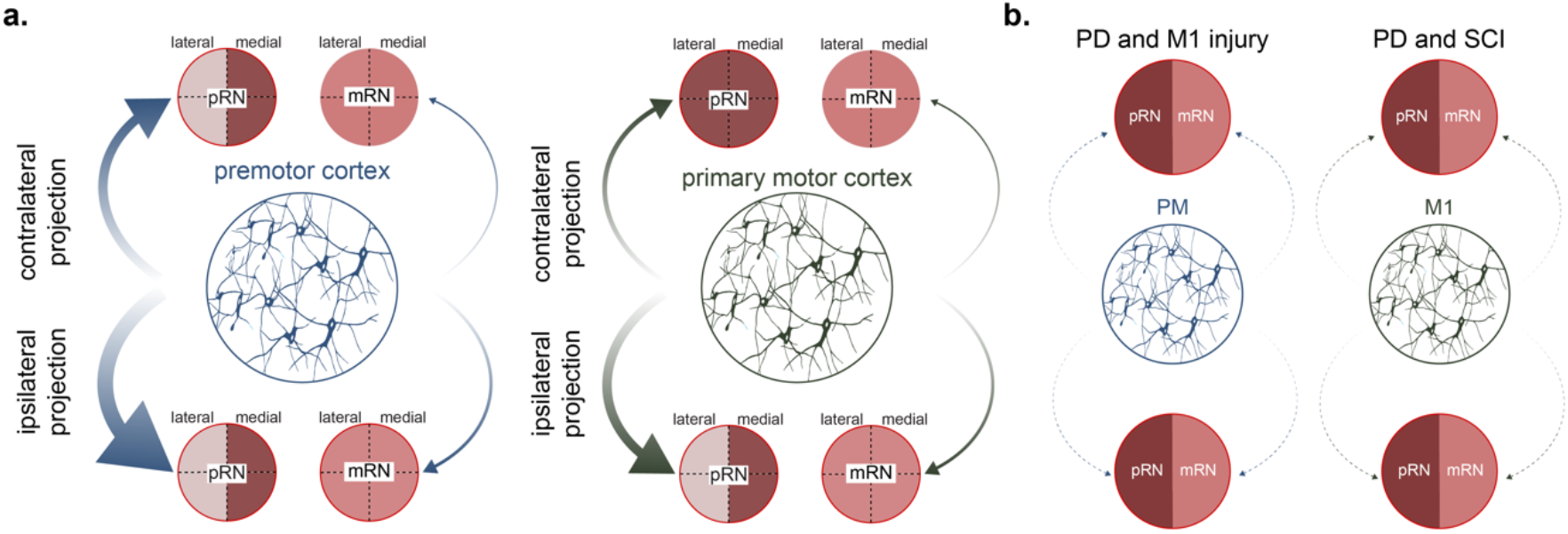
Schematic summary of the corticorubral projection density originating from PM or M1. **a.**PM corticorubral projection density is shown with blue arrows, whereas the M1 corticorubral projection density is shown with the green arrows. The larger the arrow thickness, the denser the projection. The medial preference is shown with darker red color. **b.**The corticorubral projections after multiple motor injury/disorder is drastically reduced (dashed arrows).

Although Burman and colleagues mapped the terminal boutons in the RN after injecting an anterograde tracer either in the supplementary motor area (SMA) or M1 (hand/arm regions), they could not reliably compare the axonal bouton density originating from SMA or M1 ^31^. In the present study, we implemented a semi-automated algorithm able to tackle this problematic. The algorithm detected axonal boutons within pRN and mRN and computed their respective density as well as their medio-lateral and dorso-ventral topography. First, our results are consistent with previous findings: PM and M1 mostly send their projections to the ipsilateral pRN. These projections preferentially targeted the medial portion of the pRN reflecting the somatotopical organization of the motor corticorubral projections ^14,31,33,34^. Second, our topographical analysis revealed that the projections from both PM and M1 overlapped in the medial portion of the pRN. Such convergence of motor cortical projections was also seen in the hyperdirect pathway to the STN, where PM and SMA overlap their terminal in the medial part of the STN. This convergence has been proposed to reflect the internal integration of different set of motor signals ^35^. We thus propose that pRN is capable of internally segregating PM and M1 information, integrating them and relaying the motor signal to the rubro-olivo-cerebellar tract. Finally, our quantification analysis revealed that only a small proportion of the corticorubral projections reached mRN as compared to pRN (about 20x less). This result suggests that mRN and its rubrospinal projection system might work more independently from the motor cortical areas than pRN and its rubro-olivary projection system. This hypothesis is in line with the idea that the rubrospinal system in quadrupedal animals is involved in the execution of automated movements ^36–38^. The motor cortical areas encode automated movements to a lesser extent as compared to voluntary movements ^39^. In this regard, one could expect that mRN may therefore receive less motor commands from the cerebral cortex. Interestingly, in primate, the rubrospinal system contributes to the execution of skilled hand movements ^13,28,40–43^. Our data therefore suggest that the primate mRN may act as a primary generator of skilled hand movements autonomously from the motor cortical areas ^29^. On the other hand, the motor cortical areas may contribute through pRN for sophisticated motor processing such as the acquisition and execution of conditioned reflexes ^44–47^ and motor learning through error information encoding ^48^. However, most of these studies have been conducted in non-primate animal models (cat, rabbit), where the pRN is poorly segregated in term of connectivity and morphology, as compared to primates ^10^. Future experiments are needed to further investigate the precise function of the corticorubral system in non-human primates, possibly based on reversible and selective inactivation of this projection system.

The present study provides new evidence that the projection from PM to pRN is denser than that from M1 to pRN, obtained after normalizing of the number of axonal boutons in RN by the number of corticospinal axons originating from the same PM and M1 territories injected with BDA. A similar difference between PM and M1 was found for the corticosubthalamic projection ^5^, the corticotectal projection ^6^ and the corticoreticular projection ^7^. To generalize, it seems that PM is in a position to exert a stronger influence on subcortical structures (pRN, STN, ponto-medullary reticular formation, superior colliculus) than M1. We propose that this difference in projection density to subcortical structures is related to the intrinsic function of both PM and M1. Early electrophysiological studies demonstrated that PM is responsible for both movement preparation and execution ^1,49,50^, whereas M1 is more constrained to movement execution ^2,51–58^. Indeed, recent reports showed that PM neural activity at the population level displays increased variability across motor tasks as compared to M1 ^59^. At the neural population level, Perich and colleagues demonstrated that PM operates in a higher dimension fashion than M1, likely reflecting an intrinsic neural mechanism for rapid motor learning ^60^. PM neural signatures seem therefore to exhibit more complex pattern of activity than M1. We proposed that these complex motor signals must be conveyed to subcortical structures that will in turn integrate them and transmit them back to the cortex via the thalamus. In that manner PM provides a neural substrate of advanced motor signals such as the transition between planning and motor execution ^61^.

The plastic changes in corticofugal projections from PM and M1 after motor injury or disorder differ depending on the lesion type, the lesion severity, and the respective tentative treatment ^5,8,9^ A lesion (or disorder) challenges the motor system requiring an adaptation to reestablish the best possible motor control. This adaptation can be achieved by remodeling projections (e.g. sprouting). To the best of our knowledges, we showed here for the first time a pathway (motor corticorubral projection) that undergoes the same (uniform) plastic change after multiple distinct motor injuries/disorders in primates, namely a strong decrease (nearly total loss) of the corticorubral projection from PM or M1. Therefore this uniform lesion/disorder related adaptation of the corticorubral projection differs from other descending motor projections from PM and M1 previously observed in the same groups of monkeys (see introduction and references ^5,8,9^). This unique plasticity might be a consequence of the ‘primitive’ structure of the RN, allowing the RN to play an important role after a disorder (injury) through the rubrospinal system ^62^ in synergy with the rubro-olivary system ^38^ but independently from the motor cortical areas. However, in a diffusion tensor imaging (DTI) study in chronic stroke patients, it has been shown that the alternate motor fibers (aMF), corresponding to the non-pyramidal tract fibers, displayed high fractional anisotropy (FA) in the vicinity of the red nucleus ^19^. The authors suggested that the motor corticorubral projections play a compensatory role contributing to functional recovery. Our data are not in line with this study, a discrepancy which can be explained as follows. Rüber and colleagues could not distinguish between the corticorubral and corticoreticular tracts, since both are aMF. As a result, the recovery seen by Rüber and colleagues might be due to an upregulated corticoreticular projection, observed in monkeys following sensorimotor cortex injury ^63^. Nevertheless, the primate rubrospinal neurons undergo a plastic change to facilitate their flexors and extensor muscles effects after corticospinal damage ^64^. Therefore, we suggest that the plastic changes within the RN is mediated through another mechanism than through an upregulated corticorubral projection remodeling, at least in primates. In contrast, in rats, Ishida and colleagues recently showed a causal link between an adaptative axonal sprouting of the motor corticorubral pathway and functional recovery after stroke ^65^. The discrepancy between our data and theirs may be the results of the striking difference between the respective motor system organization in rodents and in primates ^3,66,67^. Along this line, our present observation of a dramatic reduction of the corticorubral projection from PM and M1 following various motor injuries/disorders may represent an attempt to privilege direct access to the final output motor system at spinal cord level (via the direct corticospinal projection) by reducing descending collateralization to brainstem motor structures (red nucleus here), thus emphasizing the evolutionary difference recently reporting between rodents and primates ^67^.

## Materials and Methods

### Animals and BDA surgical procedures

All surgical experimental procedures, experiments, and animal care were conducted in respect to the ethical guidelines (ISBN 0-309-05377-3, 1996) and authorized by the local (Canton of Fribourg) and federal (Switzerland) veterinary authorities (authorizations No. FR 44_92_3; FR 156_00; FR 156_02; FR 156_04; FR_156_06; 156_08E; FR_157_03; FR_157_04; FR_157e_04; FR_185_08; 17_09_FR; 2012_01-FR; 2012_01E_FR). The conditions of housing in the animal facility were described earlier in detail (^24^; see also: https://tube.switch.ch/videos/2d5bb2d6). The materials and methods are similar to those reported in recent publications related to the corticobulbar (corticoreticular) ^7,8^, corticotectal projections ^6,9^ and hyperdirect corticosubthalamic pathway ^5^. Briefly, surgeries were performed under sterile conditions in anesthetized adult macaque monkeys. The protocol consisted of multiple injections of the anterograde tracer BDA (MW = 10,000; Molecular Probe, Eugene, OR, USA) unilaterally in PM or M1, using a 10μl Hamilton micro-syringe. BDA concentration was 5% (in distilled water) for M9380 and 10% for the other monkeys. The extent of the BDA injection site was assessed on consecutive histological frontal sections, each reconstructed manually using Neurolucida software (MBF Bioscience-MicroBrightField Inc., version 11). Three-dimension (3D) reconstruction was performed by stacking on a dorsolateral view the medio-lateral extent of BDA intake from the midline. The injection sites in PM involved both PMd and PMv in most monkeys, except in R12, LL and RO in which BDA was delivered mostly in PMd. The BDA injection in M1 was limited to the hand area in M9380, as defined by intracortical microstimulation. For the other M1 injections, based on anatomical landmarks (cortical sulci), BDA involved the hand area, as well as more proximal territories of the forelimb. All the detailed information (e.g. monkey age, BDA volume, etc) have been previously published ^5–9^.

### Lesion and treatment procedures

The first lesion group was composed by six monkeys (CS, CP, CG, AC, AG and AP) that underwent a cervical spinal cord injury (SCI) at C7-C8 level (hemisection). This SCI group was further subdivided into a control subgroup (n=3; CS, CP and CG) and subgroup subjected to an anti-Nogo-A antibody treatment (n=3; AC, AG and AP; see ^20–22^). The surgical, lesion and behavioral procedures have been previously reported in ^21–23^. Briefly, the six monkeys were treated with antibodies (control or anti-Nogo-A) applied intrathecally by infusion over 4 weeks into the subdural space of the lower cervical spinal cord. In both subgroups, the treatment was delivered with an osmotic pump (Alzet, 2ML2, 2mL) located in the animal’s back 3–5 mm rostral to the cervical lesion site and connected to it by a silastic tube. The implantation of the osmotic pump was performed a few minutes after the cervical lesion itself.

The surgical, lesion and behavioral procedures of the group of 4 monkeys (LL, MY, LY and MI) subjected to Parkinson’s disease-like symptoms (PD) have been previously reported in ^24,25,68^. Briefly, the four monkeys underwent daily MPTP (0.5 mg/kg, Sigma; i.m.; duration: about 1 month) injections with the purpose to kill dopaminergic neurons in the substantia nigra pars compacta (SNpc). At the end of the lesional protocol the monkeys had received between 6.25 ml/ kg and 7.75 mg/kg of MPTP. The protocol consisted of two series of injections for 4 days each with a break after the fourth day. All safety protocols were adapted in accordance to the American national institute of health (NIH). After a stable plateau of incomplete spontaneous functional recovery was reached, the four monkeys were treated with autologous neural cell ecosystems (ANCE) obtained by cell culture of cortical neurons developed in vitro from cortical biopsies ^24,25,68–74^. ANCE were then reimplanted in the same monkey. The injections sites (two in the Putamen and one in the Caudate nucleus on each side) were identified by aid of postlesional T1-weighted MRI scan (OsiriX, v 4.1.2) and confirmed by the Paxinos atlas ^75^. Cells were implanted via a Hamilton microsyringe (10μl culture medium; 2μl/ min during 5 min per injection site) using a nanoinjector (Stoelting, Wood Dale, IL, USA) fixed to a stereotaxic frame. Each monkey received between 250,000 and 400,000 cells. At the end of the protocol, two of the PD monkeys were injected unilaterally with BDA in PM (LL, MY) and the other two monkeys in M1 (LY, MI).

Finally, a group (MCI) of four monkeys, subjected to a M1-injury (hand area), were injected in PM with BDA unilaterally. Similar to the SCI group, two MCI monkeys received an anti-Nogo-A antibody treatment and two MCI monkeys were untreated (controls). The four M1 cortically lesioned monkeys (RO, BI, VA and MO) were implanted with chronic chambers fixed to the skull for easier access to the hand region of M1. Cortical lesion of M1 hand area was obtained by multiple injections of ibotenic acid (10 μg/μl in phosphate buffered saline) with a 10μl Hamilton syringe at multiple sites corresponding to digits areas previously identified by ICMS, as previously reported ^26,27^. VA and MO were treated with anti-Nogo-A antibodies (3mg/mL) administrated by two osmotic pumps (Alzet, model 2ML2, 5μl/hr) implanted under anesthesia in a subcutaneous pouch in the neck region. One pump administered the treatment intrathecally to the cervical spinal cord, whereas the other pump delivered the antibody close to the lesion site in M1 below the dura. The treatment was delivered for a total duration of 4 weeks before surgical removal of the two osmotic pumps. The treatment was delivered immediately after the lesion, thus the surgery for pumps implantation followed the injection of ibotenic acid.

### Histological procedures

At the end of the experimental protocols, after completion of behavioral and electrophysiological investigations, BDA injections and survival time to allow axonal transport of BDA, the monkeys were euthanized under lethal anesthesia. Surgical procedures for sacrifice were previously reported ^21–24,26,27^. Briefly, the monkeys were subjected to transcardiac perfusion with 400 ml, 0.9% saline, followed by 3 Liters of 4% paraformaldehyde (0.1 M phosphate buffer, pH 7.6) and finally three series of 2 Liters each of sucrose solutions (10%, 20% and 30%) to fix and cryopreserve the tissue. The brains were cut in the frontal plane in 50μm thick sections with a freezing microtome, collected in 5–7 series. One series of brain sections was processed to visualize BDA in order to reconstruct the location of stem axons and axonal boutons, both *en passant* and *terminaux* (histochemical protocol described in ^76^). Another series of sections was used to stain for Nissl bodies in order to visualize the cytoarchitecture of the red nucleus.

### Data collection and analysis

To quantify the synaptic terminal and *en passant* axonal boutons within the RN, we developed an algorithm that detects BDA-stained boutons. We imaged the histological sections (Nissl and BDA-stained) with the bright field microscope slide scanner Leica DM6B Navigator, the camera DMC5400 and LAS X Navigator software at magnification of 10x. We analyzed all the histological sections within the 2 Nissl and BDA series covering the RN in the intact monkeys. Due to the low (or inexistent) number of axonal boutons in the injured monkeys (SCI, MCI, PD), we analyzed only sections where we found axonal boutons. If a monkey did not show any axonal boutons, we performed the analysis in four sections only. For all the subsequent analysis we used MATLAB R2019a. First, the images were converted into jpeg format from their original lif format. Then, we manually co-registered the two corresponding BDA and Nissl-stained sections and defined the pRN and mRN contours on the resulting images. Based on the contours, we extracted the pRN and mRN and converted the images in grayscale. We then converted the grayscale images into binary images by applying a threshold of gray pixel intensity. Different thresholds were tested on few random sections for each monkey. We choose the optimal threshold by visually verifying that the resulting binary image captured actual axonal boutons. Once the threshold defined, we applied the same threshold for all the sections in each monkey and obtained the binary images. Since BDA protocol may result in staining artifacts (e.g. blood vessels and neurons), we applied two filters: (i) a size filter of conglomerated pixels with a lower and upper size limit of 2 and 400 pixels, respectively; and (ii) a circularity, *C*, filter of conglomerated pixels of 0.5 (1 being a ‘perfect’ circle) based on the following formula:

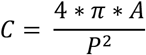

where *C* is the circularity, *A* is the area of the conglomerated pixels and *P* is the perimeter of the conglomerated pixels. These two parameters were kept for all histological sections in all monkeys. We tested the accuracy and validity of the algorithm by comparing in one intact monkey (CH) the numbers of axonal boutons obtained either with the present algorithm or by manually charting all axonal boutons using the manual exhaustive plotting method in RN as we did previously in recent studies on the corticoreticular, corticotectal and corticosubthalamic projections (5-9). We obtained consistent results with the 2 methods, attested by comparable distributions of axonal boutons along the rostrocaudal axis of RN in monkey CH. The present automated and the previous manual ^5,8,9^ counting methods both consist of an exhaustive sampling approach of axonal boutons in a given target nucleus, for which stereology is not pertinent. We then normalized in each monkey the number of axonal boutons in RN with the number of corticospinal axons labeled just above the pyramidal decussation as described in previous reports ^5–9^. The axonal bouton density of each section was obtained by counting the number of axonal boutons in a 2,500μm^2^ area. For each histological section, we divided pRN and mRN into four quadrants: dorso-lateral, dorso-medial, ventral-lateral and ventro-medial parts. To compare the density topography across sections, we interpolated (MATLAB function *interp1*) the density of each quadrant into equal sizes and plotted the average density across sections. The density of each section was measured with the area under the curves (AUC, MATLAB function *trapz*). Due to the low number of monkeys, we performed a bootstrapping analysis, where the initial population of n analyzed histological sections, covering the rostrocaudal extent of the pRN or mRN, was resampled with replacement to obtain k = 100,000 fictive populations of n sections independently for each monkey. For each bootstrapped population, we summed up the number of BDA-labeled boutons along the rostrocaudal axis to finally obtain the total number of BDA-labeled boutons of the 100,000 bootstrapped populations.

As standard in bootstrapping, we calculated the p-value by estimating the residuals of the combined distribution testing for the null hypothesis that the gaussian distribution of the difference had a mean value of 0:

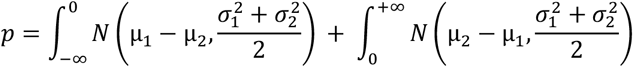

where *N* is the gaussian distribution function; μ_1_ and μ_2_ are the means of the two conditions with μ_1_>μ_2_ and *σ*_1_ and *σ*_2_ are their respective standard deviations. Finally, the significance threshold of the p-value has been adjusted using the Bonferroni correction for multiple-group comparisons (α = 0.05 divided by the number of groups). This bootstrapping methodology has been previously performed in another report ^5^. Some analyses did not require a bootstrapping analysis (e.g. the comparison between number of axonal boutons between pRN and mRN). We therefore used the standard Wilcoxon signed rank test. The performed statistical tests as well as the α thresholds after Bonferroni correction are mentioned in the captions of each plot.

## Acknowledgments

We thank Prof. Joceylne Bloch, Prof. Patrick Freund, Dr. Jean-Francois Brunet, Dr. Simon Badoud and Dr. Eric Schmidlin for surgical assistance; Prof. Martin Schwab, Prof. Joceylne Bloch, Prof. Patrick Freund, Dr. Anis Mir, Dr. T. Wannier, Dr. Eric Schmidlin, Dr. Marie-Laure Beaud, Dr. Alexander Wyss, Dr. Shahid Bashir, Dr. Adjia Hamadjida, Dr. Aderraouf Belhaj-Saif, Dr. Mélanie Kaeser, Dr. J. Savidan, Dr. Anne-Dominique Gindrat, Dr. Yu Liu, Dr. Michela Fregosi, Dr. Simon Badoud, Mr. Alessandro Contestabile and Mr. Jérôme Cottet for contributions to experimental sessions and elaborations of experimental protocols; Mr. L. Bossy, Mr. J. Maillard, Mr. B. Bapst, Mr. B. Morandi and Mr. J. Corpataux for animal care assistance; Mrs. Christine Roulin, Mrs. Véronique Moret and Mrs. Christiane Marti for tissue processing. Mr. Felix Meyenhofer for technical microscopical assistance. Finally, we thank Dr. Erinn M. Grigsby for the proof-reading. This work was financially supported by Swiss National Science Foundation Grants No 110005, 132465, 144990, 149643, Sinergia CRSII3_160696, Sinergia PROMETHEUS CRSI33_125408 (to EMR), postdoc mobility No 210986 (to SB) and the Swiss Primate Competence Centre for Research (SPCCR: www.unifr.ch/spccr).

## Author Contributions

SB and EMR designed the research project; SB and EMR performed the research; SB analyzed the data; SB and EMR wrote the paper.

## Supplementary figures

**Supplementary Figure 1.**
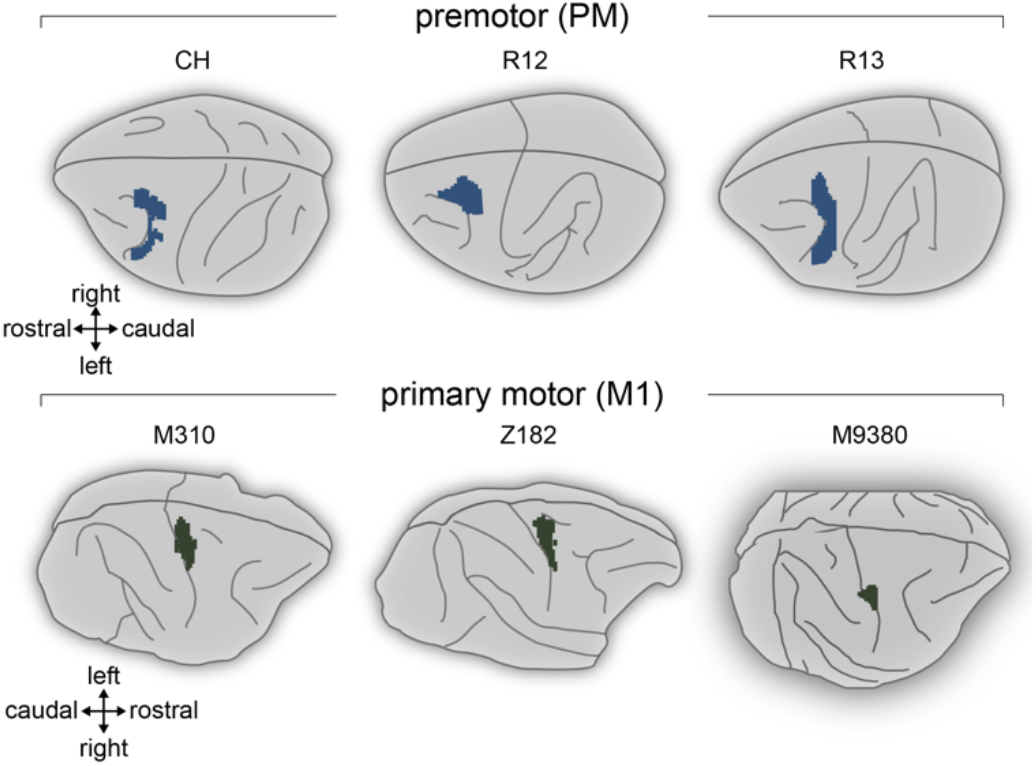
Reconstruction of the intact monkeys’ brain injected with BDA in the premotor cortex (PM) or in the primary motor cortex (M1). The extent of BDA uptake was calculated on each histological frontal section and were transposed to the 3D brain (see ^5^). Redrawn from previous publications ^5–9^.

**Supplementary Figure 2.**
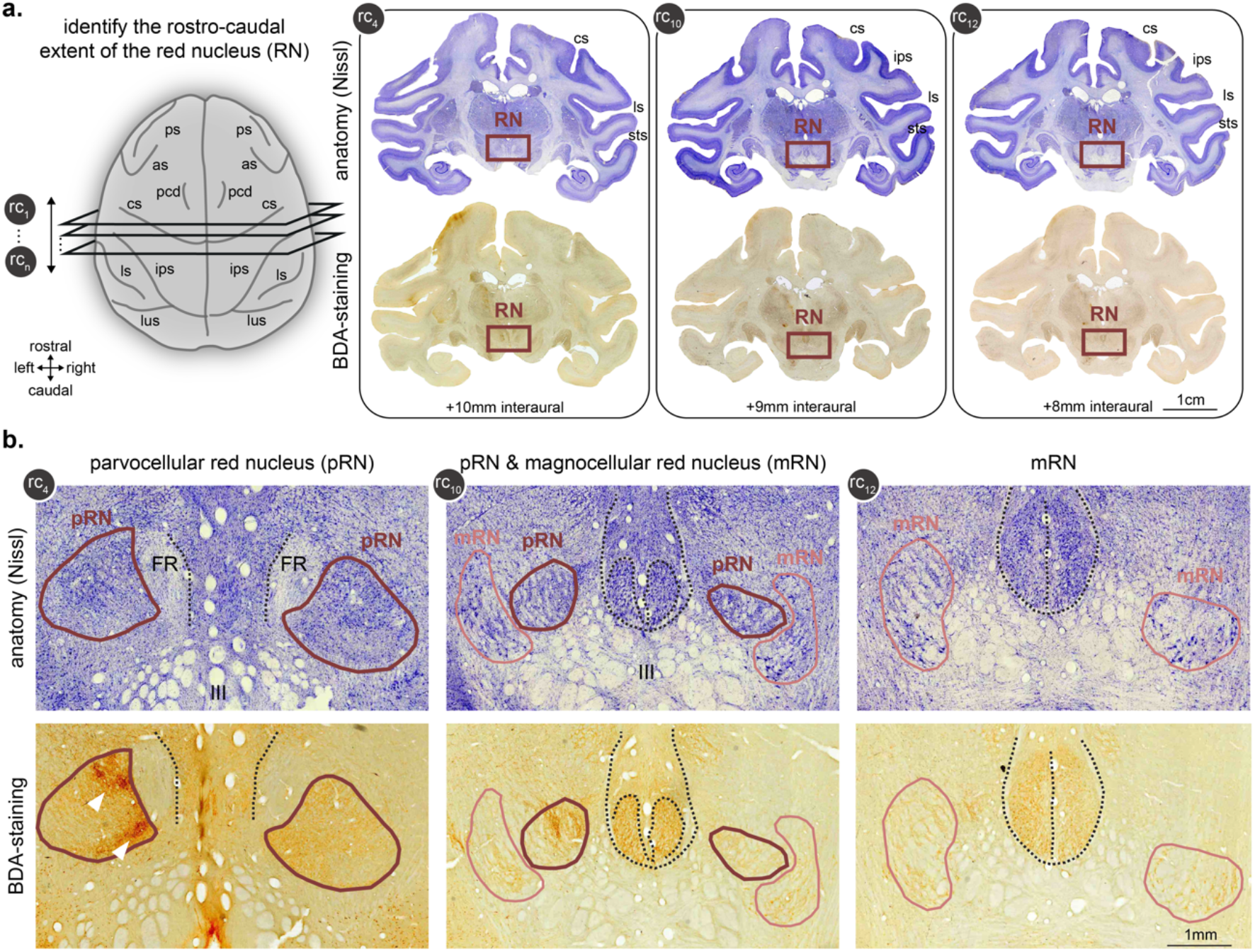
Identification of the red nucleus on frontal histological sections. **a.**Examples of 3 histological sections (rc4, rc10 and rc12) in a representative monkey (CH). Top row: we identify the red nucleus on Nissl-stained section. Bottom row: the photographs show the corresponding BDA-staining sections. **b.**Magnification of the 3 histological sections. Top row: In the histological section rc4, the parvocellular red nucleus (pRN) is visible (dark red contour). In the histological section rc10, the pRN (dark red contour) and the magnocellular red nucleus (mRN) are visible (light red contour). In the histological section rc12, only the mRN is visible (light red contour). Bottom row: the RN corresponding coutours are drawn in the BDA-stained sections. Note the dense corticorubral projection in section rc4 (white arrows), whereas rc10 shows less BDA-stained signal and rc12 has no signal. FR: fasciculus retroflexus, III: oculomotor nerve. Dashed lines represent the periaqueductal gray matter.

**Supplementary Figure 3.**
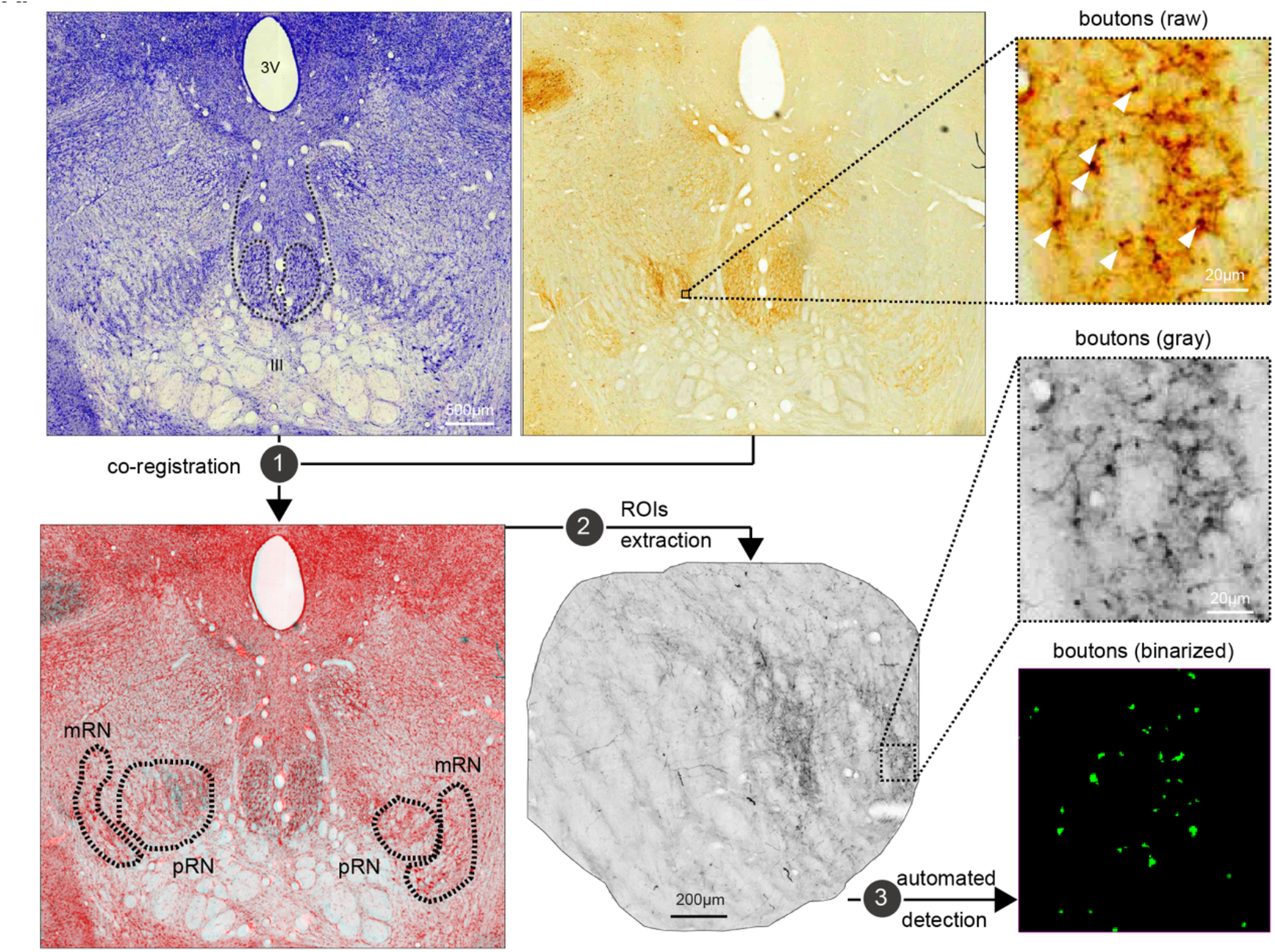
Methodology of axonal corticorubral boutons detection in RN. The example shows the histological section rc10 of **Supplementary Figure 2**. Step 1: the two corresponding histological sections (Nissl-stained and BDA-stained sections) were manually co-registered. In the resulting co-registered image, we manually drew the contours (dashed lines) of the left pRN, left mRN, right pRN and right mRN. Step 2: each contour resulted in a gray colored ROI extraction of the BDA-stained section (here the example is the left (ipsilateral) pRN. Step 3: we implemented an algorithm that detected the axonal boutons based on the intensity, the circularity and the size of conglomerated pixels; and resulted in a binarized axonal boutons image.

**Supplementary Figure 4.**
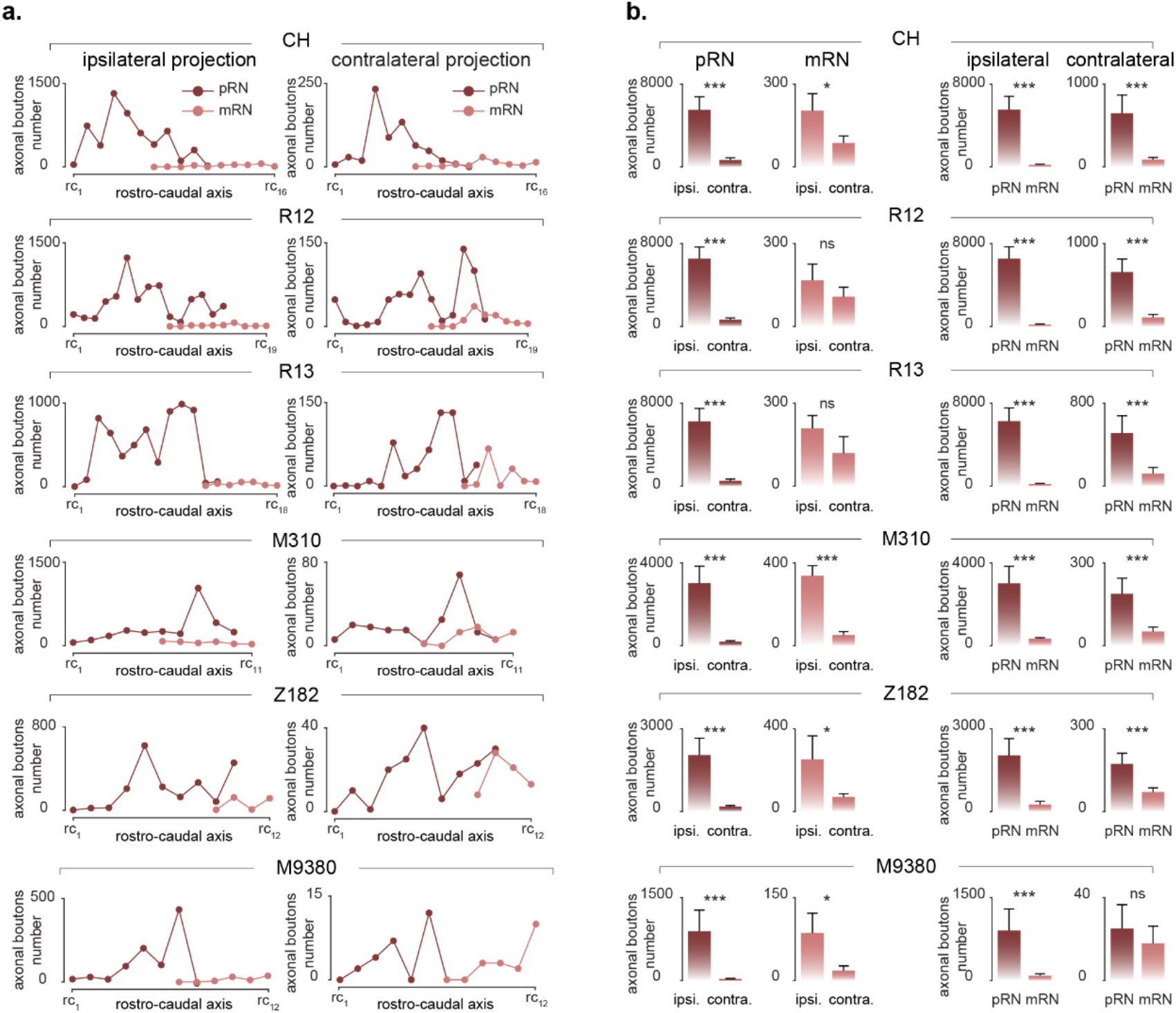
Corticorubral axonal boutons numbers for each individual intact monkey. **a.** Plots showing the axonal boutons numbers in all the analysed histological sections spanning the red nucleus. The histological sections are organized following the rostro-caudal axis (rc) with rc1 being the most rostral section where pRN is visible and rc_n_ being the most caudal section where mRN is visible. **b.** Bar plots showing the means + the standard deviations of axonal boutons numbers for each individual monkey across bootstrapped histological sections (see Methods). All statistically significances are shown with * for p-values < 0.05, ** for p-values < 0.01, *** for p-values < 0.001, “ns” meaning statistically non-significant. P-values are calculated with the residual estimation of the combined distributions.

**Supplementary Figure 5.**
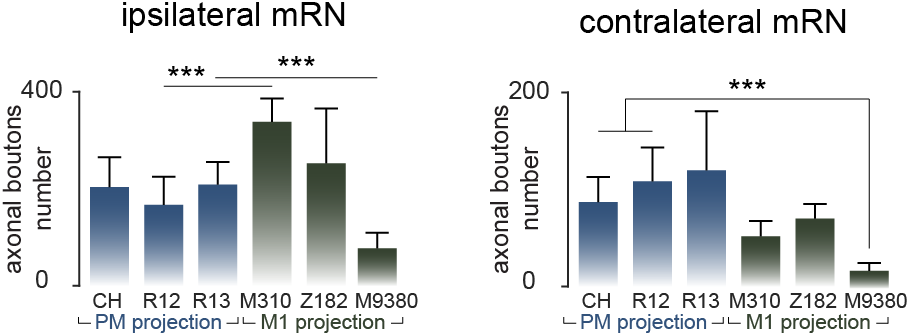
Corticorubral projection density in mRN in the 6 intact monkeys. **a.** Bar plots showing the means + the standard deviations of axonal boutons numbers for each individual intact monkey across bootstrapped histological sections (see Methods). All statistically significances are shown with *** for p-values < 0.00016, residual estimation of the combined distributions and Bonferroni correction for multiple-group comparisons.

**Supplementary Figure 6.**
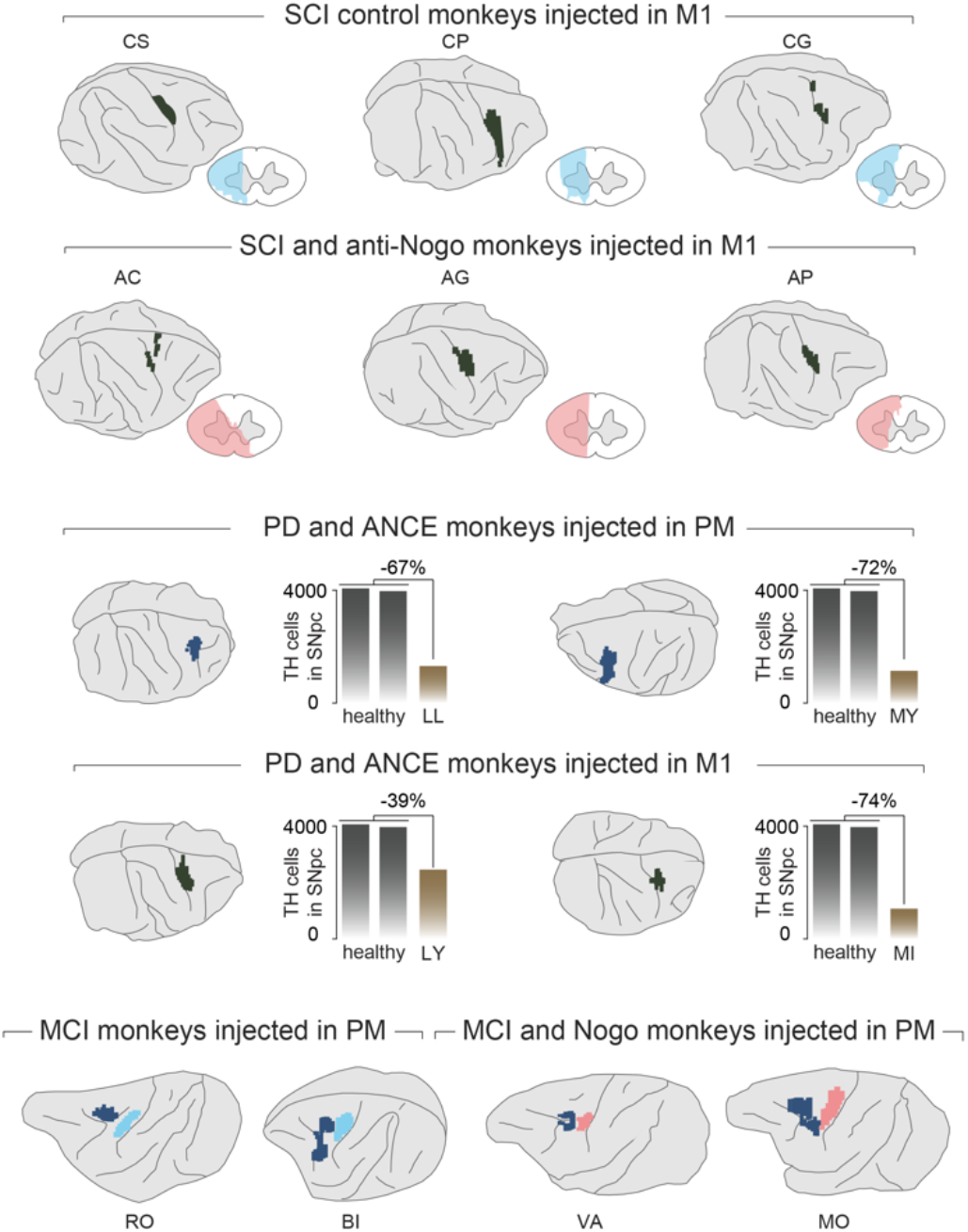
Reconstruction of the lesioned monkeys injected unilaterally with BDA in the premotor cortex (PM) or in the primary motor cortex (M1). The extent of BDA uptake was calculated on each histological frontal section and were transposed to the 3D brain (see ^5^). The spinal cord lesion extents (C7-C8 level) are redrawn from ^21–23^. The thyrosine-hydroxylase (TH) neurons in the substantia nigra pars compacta (SNpc) were used to measure the degree of dopamine loss after Parkinson’s disease-like symptoms. This analysis has been previously shown in ^24^. The primary motor cortex (M1) lesion extent was measured on frontal histological sections and transposed on the 3D brain of each monkey. The lesion and BDA uptake reconstructions were redrawn from ^26,27^.

